# An infectious SARS-CoV-2 B.1.1.529 Omicron virus escapes neutralization by several therapeutic monoclonal antibodies

**DOI:** 10.1101/2021.12.15.472828

**Authors:** Laura A. VanBlargan, John M. Errico, Peter J. Halfmann, Seth J. Zost, James E. Crowe, Lisa A. Purcell, Yoshihiro Kawaoka, Davide Corti, Daved H. Fremont, Michael S. Diamond

**Author notes:** **Corresponding author:** Michael S. Diamond, M.D., Ph.D.

## Abstract

Severe acute respiratory syndrome coronavirus 2 (SARS-CoV-2) has caused the global COVID-19 pandemic resulting in millions of deaths worldwide. Despite the development and deployment of highly effective antibody and vaccine countermeasures, rapidly-spreading SARS-CoV-2 variants with mutations at key antigenic sites in the spike protein jeopardize their efficacy. Indeed, the recent emergence of the highly-transmissible B.1.1.529 Omicron variant is especially concerning because of the number of mutations, deletions, and insertions in the spike protein. Here, using a panel of anti-receptor binding domain (RBD) monoclonal antibodies (mAbs) corresponding to those with emergency use authorization (EUA) or in advanced clinical development by Vir Biotechnology (S309, the parent mAbs of VIR-7381), AstraZeneca (COV2-2196 and COV2-2130, the parent mAbs of AZD8895 and AZD1061), Regeneron (REGN10933 and REGN10987), Lilly (LY-CoV555 and LY-CoV016), and Celltrion (CT-P59), we report the impact on neutralization of a prevailing, infectious B.1.1.529 Omicron isolate compared to a historical WA1/2020 D614G strain. Several highly neutralizing mAbs (LY-CoV555, LY-CoV016, REGN10933, REGN10987, and CT-P59) completely lost inhibitory activity against B.1.1.529 virus in both Vero-TMPRSS2 and Vero-hACE2-TMPRSS2 cells, whereas others were reduced (∼12-fold decrease, COV2-2196 and COV2-2130 combination) or minimally affected (S309). Our results suggest that several, but not all, of the antibody products in clinical use will lose efficacy against the B.1.1.529 Omicron variant and related strains.

## MAIN TEXT

Since December of 2019, the global COVID-19 pandemic caused by SARS-CoV-2 has resulted in 267 million infections and 5.3 million deaths. The expansion of the COVID-19 pandemic and its accompanying morbidity, mortality, and destabilizing socioeconomic effects have made the development and distribution of SARS-CoV-2 therapeutics and vaccines an urgent global health priority^1^. While the rapid deployment of countermeasures including monoclonal antibodies and multiple highly effective vaccines has provided hope for curtailing disease and ending the pandemic, this has been jeopardized by emergence of more transmissible variants with mutations in the spike protein that also could evade protective immune responses.

Indeed, over the past year, several variant strains have emerged including B.1.1.7 (Alpha), B.1.351 (Beta), B.1.1.28 [also called P.1, Gamma]), and B.1.617.2 (Delta), among others, each having varying numbers of substitutions in the N-terminal domain (NTD) and the RBD of the SARS-CoV-2 spike. Cell-based assays with pseudoviruses or authentic SARS-CoV-2 strains suggest that neutralization by many EUA mAbs might be diminished against some of these variants, especially those containing mutations at positions L452, K477, and E484^2-6^. Notwithstanding this, *in vivo* studies in animals showed that when most EUA mAbs were used in combination they retained efficacy against different variants^7^. The recent emergence of B.1.1.529, the Omicron variant^8,9^, which has a larger number of mutations (∼30 substitutions, deletions, or insertions) in the spike protein, has raised concerns that this variant will escape from protection conferred by vaccines and therapeutic mAbs.

We obtained an infectious clinical isolate of B.1.1.529 from a symptomatic individual in the United States (hCoV-19/USA/WI-WSLH-221686/2021). We propagated the virus once in Vero cells expressing transmembrane protease serine 2 (TMPRSS2) to prevent the emergence of adventitious mutations at or near the furin cleavage site in the spike protein^10^. Our B.1.1.529 isolate encodes the following mutations in the spike protein (A67V, Δ69−70, T95I, G142D, Δ143-145, Δ211, L212I, insertion 214EPE, G339D, S371L, S373P, S375F, K417N, N440K, G446S, S477N, T478K, E484A, Q493R, G496S, Q498R, N501Y, Y505H, T547K, D614G, H655Y, N679K, P681H, N764K, D796Y, N856K, Q954H, N969K, and L981F; **Fig 1a-b** and GISAID: EPI_ISL_7263803), which is similar to strains identified in Africa^11^. Our isolate, however, lacks an R346K mutation, which is present in a minority (∼8%) of reported strains.

**Figure 1.**
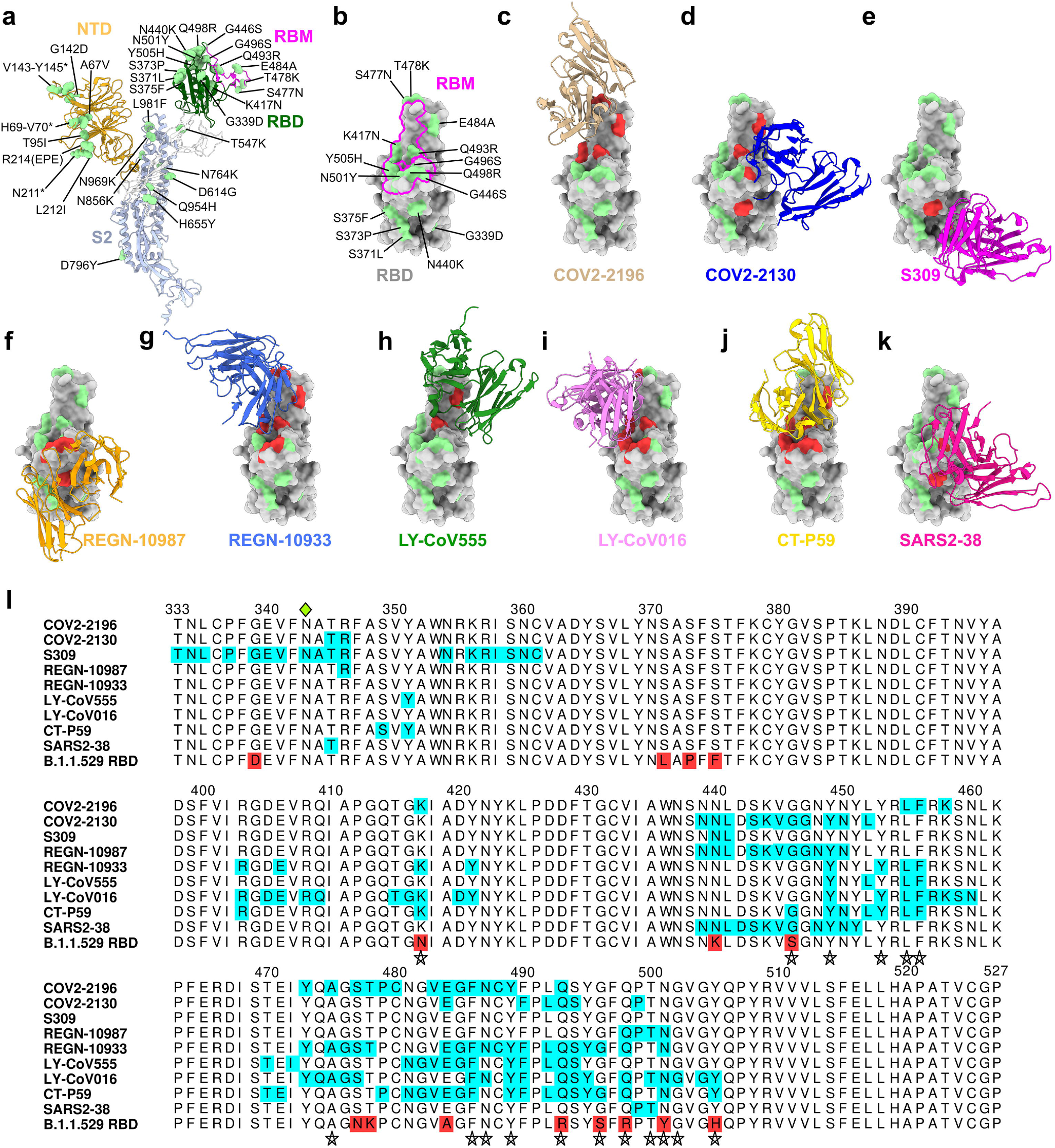
Neutralizing mAb epitopes on B.1.1.529. **a-b**, SARS-CoV-2 spike trimer (PDB: 7C2L and PDB: 6W41). One spike protomer is highlighted, showing the NTD in orange, RBD in green, RBM in magenta, and S2 portion of the molecule in blue (**a**). Close-up view of the RBD with the RBM outlined in magenta (**b**). Amino acids that are changed in B.1.1.529 compared to WA1/2020 are indicated in light green (**a-b**), with the exception of N679K and P681H, which were not modeled in the structures used. **c-k**, SARS-CoV-2 RBD bound by EUA mAbs COV2-2196 (**c**, PDB: 7L7D); COV2-2130 (**d**, PDB: 7L7E); S309 (**e**, PDB: 6WPS); REGN-10987 (**f**, PDB: 6XDG); REGN-10933 (**g**, PDB: 6XDG)); LY-CoV555 (**h**, PDB: 7KMG) LY-CoV016 (**i**, PDB: 7C01); CT-P59 (**j** PDB: 7CM4) and SARS2-38 (**k**, PDB: 7MKM). Residues mutated in the B.1.1.529 RBD and contained in these mAbs respective epitopes are shaded red, whereas those outside the epitope are shaded green. **l**, multiple sequence alignment showing the epitope footprints of each EUA mAb on the SARS-CoV-2 RBD highlighted in cyan. B.1.1.529 RBD is shown in the last row, with sequence changes relative to the WT RBD highlighted red. A green diamond indicates the location of the N-linked glycan at residue 343. Stars below the alignment indicate hACE2 contact residues on the SARS-CoV-2 RBD^38^.

Given the number of substitution in the B.1.1.529 spike protein, including eight amino acid changes (K417N, G446S, S477N, Q493R, G496S, Q498R, N501Y, Y505H) in the ACE2 receptor binding motif (RBM), we first evaluated possible effects on the structurally-defined binding epitopes of mAbs corresponding to those with EUA approval or in advanced clinical development (S309 [parent of VIR-7381]^12,13^; COV2-2196 and COV2-2130 [parent mAbs of AZD8895 and AZD1061, respectively]^14^; REGN10933 and REGN10987^15^, LY-CoV555 and LY-CoV016^16,17^; and CT-P59 [Celltrion]^18^) along with an additional broadly neutralizing mAb (SARS2-38) that we recently described^19^. We mapped the B.1.1.529 spike mutations onto the antibody-bound SARS-CoV-2 spike or RBD structures published in the RCSB Protein Data Bank (**Fig 1c-k**). While every antibody analyzed had structurally defined recognition sites that were altered in the B.1.1.529 spike, the differences varied among mAbs with some showing larger numbers of changed residues (**Fig 1l**: COV2-2196, n = 4; COV2-2130, n = 4; S309, n = 2; REGN10987, n = 4; REGN10933, n = 8; Ly-CoV555, n = 2; Ly-CoV016, n = 6; CT-P59, n = 8; and SARS2-38, n = 2).

To address the functional significance of the spike sequence variation in B.1.1.529 for antibody neutralization, we used a high-throughput focus reduction neutralization test (FRNT)^20^ with WA1/2020 D614G and B.1.1.529 in Vero-TMPRSS2 cells (**Fig 2**). We tested individual and combinations of mAbs that target the RBD in Vero-TMPRSS2 cells including S309 (Vir Biotechnology), COV2-2130/COV2-2196 (parent mAbs of AZD1061 and AZD8895 provided by Vanderbilt University), REGN10933/REGN10987 (synthesized based on casirivimab and imdevimab sequences from Regeneron), LY-CoV555/LY-CoV016 (synthesized based on bamlanivimab and etesevimab sequences from Lilly), CT-P59 (synthesized based on regdanvimab sequences from Celltrion), and SARS2-38. As expected, all individual or combinations of mAbs tested neutralized the WA1/2020 D614G isolate with EC_50_ values similar to published data^6,18,21^. However, when tested alone, REGN10933, REGN10987, LY-CoV555, LV-CoV016, CT-P59 and SARS2-38 completely lost neutralizing activity against B.1.1.529, with little inhibitory capacity even at the highest (10,000 ng/mL) concentration tested. COV2-2130 and COV2-2196 showed an intermediate ∼12 to 150-fold (*P* < 0.0001) loss in inhibitory activity, respectively against the B.1.1.529 strain. In comparison, S309 showed a less than 2-fold (*P* > 0.5) reduction in neutralizing activity against B.1.1.529 (**Fig 2a-h**). Analysis of mAb combinations currently in clinical use showed that REGN10933/REGN10987 and LY-CoV555/LV-CoV016 lost all neutralizing activity against B.1.1.529, whereas COV2-2130/COV2-2196 showed a ∼12-fold (*P* < 0.0001) reduction in inhibitory activity.

**Figure 2.**
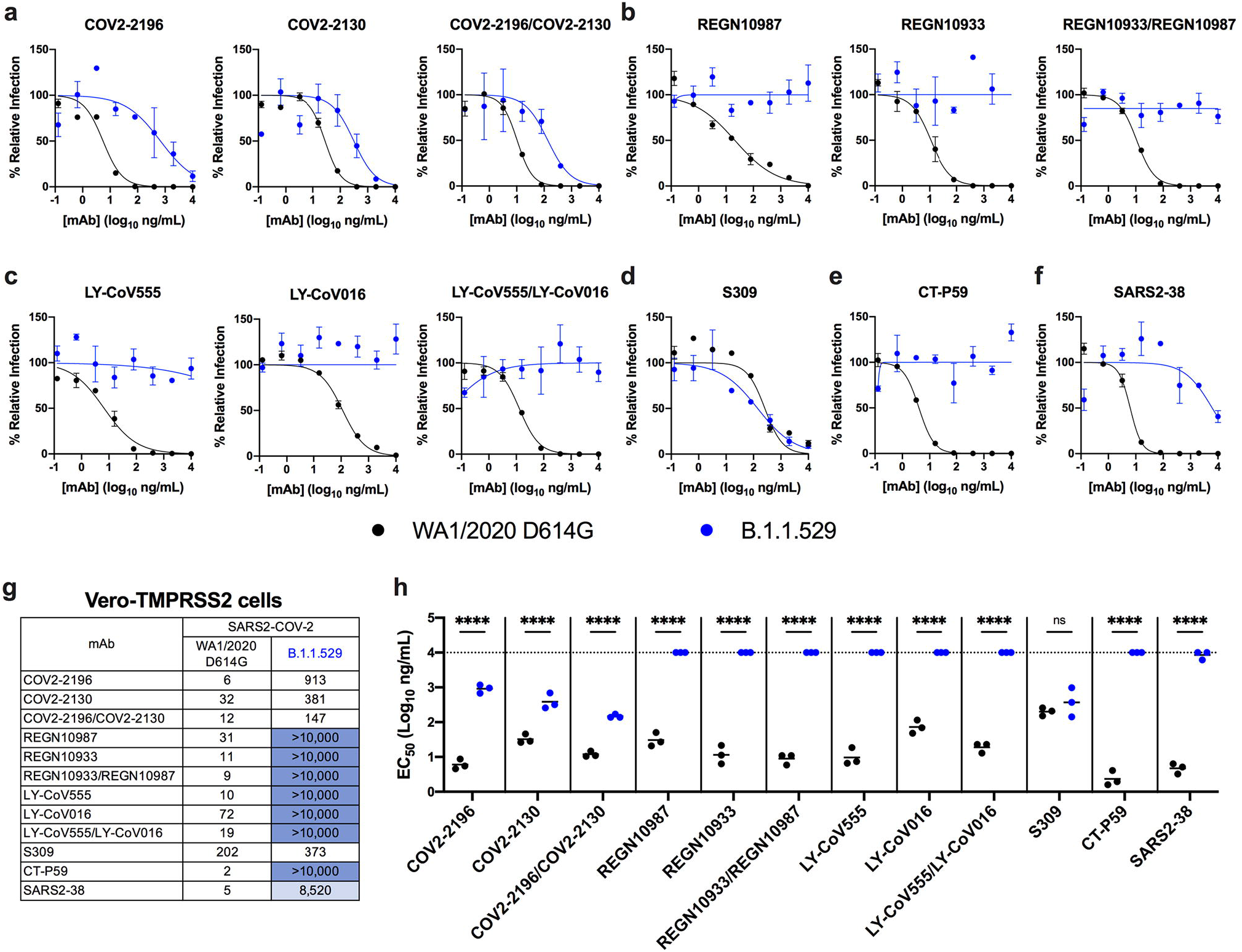
Neutralization of SARS-CoV-2 B.1.1.529 Omicron strain by mAbs in Vero-TMPRSS2 cells. **a-f**, Neutralization curves in Vero-TMPRSS2 cells comparing the sensitivity of SARS-CoV-2 strains with the indicated mAbs (COV2-2196, COV2-2130; REGN10933, REGN10987, LY-CoV555, LY-CoV016, S309, CT-P59, and SARS2-38) with WA1/2020 D614G and B.1.1.529. Also shown are the neutralization curves for antibody cocktails (COV2-2196/COV2-2130, REGN10933/REGN10987, or LY-CoV555/LY-CoV016). One representative experiment of three performed in technical duplicate is shown. Error bars indicate range. **g**, Summary of EC_50_ values (ng/ml) of neutralization of SARS-CoV-2 viruses (WA1/2020 D614G and B.1.1.529) performed in Vero-TMPRSS2 cells. Data is the geometric mean of 3 experiments. Blue shading: light, EC_50_ > 5,000 ng/mL; dark, EC_50_ > 10,000 ng/mL. **h**, Comparison of EC_50_ values by mAbs against WA1/2020 D614G and B.1.1.529 (3 experiments, ns, not significant; ^****^, *P* < 0.0001; two-way ANOVA with Sidak’s post-test). Bars indicate mean values. The dotted line indicates the upper limit of dosing of the assay.

We repeated experiments in Vero-hACE2-TMPRSS2 cells to account for effects of hACE2 expression, which can affect neutralization by some anti-SARS-CoV-2 mAbs^19,22^. Moreover, modeling studies suggest that the mutations in the B.1.1.529 spike may enhance interactions with hACE2^23^. All individual or combinations of mAbs tested neutralized the WA1/2020 D614G isolate as expected. However, REGN10933, REGN10987, LY-CoV555, LV-CoV016, SARS2-38, and CT-P59 completely lost neutralizing activity against B.1.1.529, and the combinations of REGN10933/ REGN10987 or LY-CoV555/LV-CoV016 also lacked inhibitory capacity (**Fig 3a-h**). In comparison, COV2-2196 showed moderately reduced activity (∼16-fold) as did the combination of COV2-2130/COV2-2196 mAbs (∼11-fold). Unexpectedly, COV2-2130 did not show a difference in neutralization of WA1/2020 and B.1.1.529 in the Vero-hACE2-TMPRSS2 cells (**Fig 3a, g, and h**), whereas it did in Vero-TMPRSS2 cells (**Fig 2a, g, and h**). The S309 mAb showed less potent neutralizing activity in Vero-hACE2-TMPRSS2 cells at baseline with a flatter dose response curve (**Fig 3d**), as seen previously^6,24^, and showed a moderate (∼6-fold, *P* < 0.0001) reduction in neutralizing activity against B.1.1.529 compared to WA1/2020 D614G. Thus, while the trends in mAb neutralization of B.1.1.529 generally were similar to Vero-TMPRSS2 cells, some expected and unexpected differences were noted with COV2-2130 and S309 on cells expressing hACE2.

**Figure 3.**
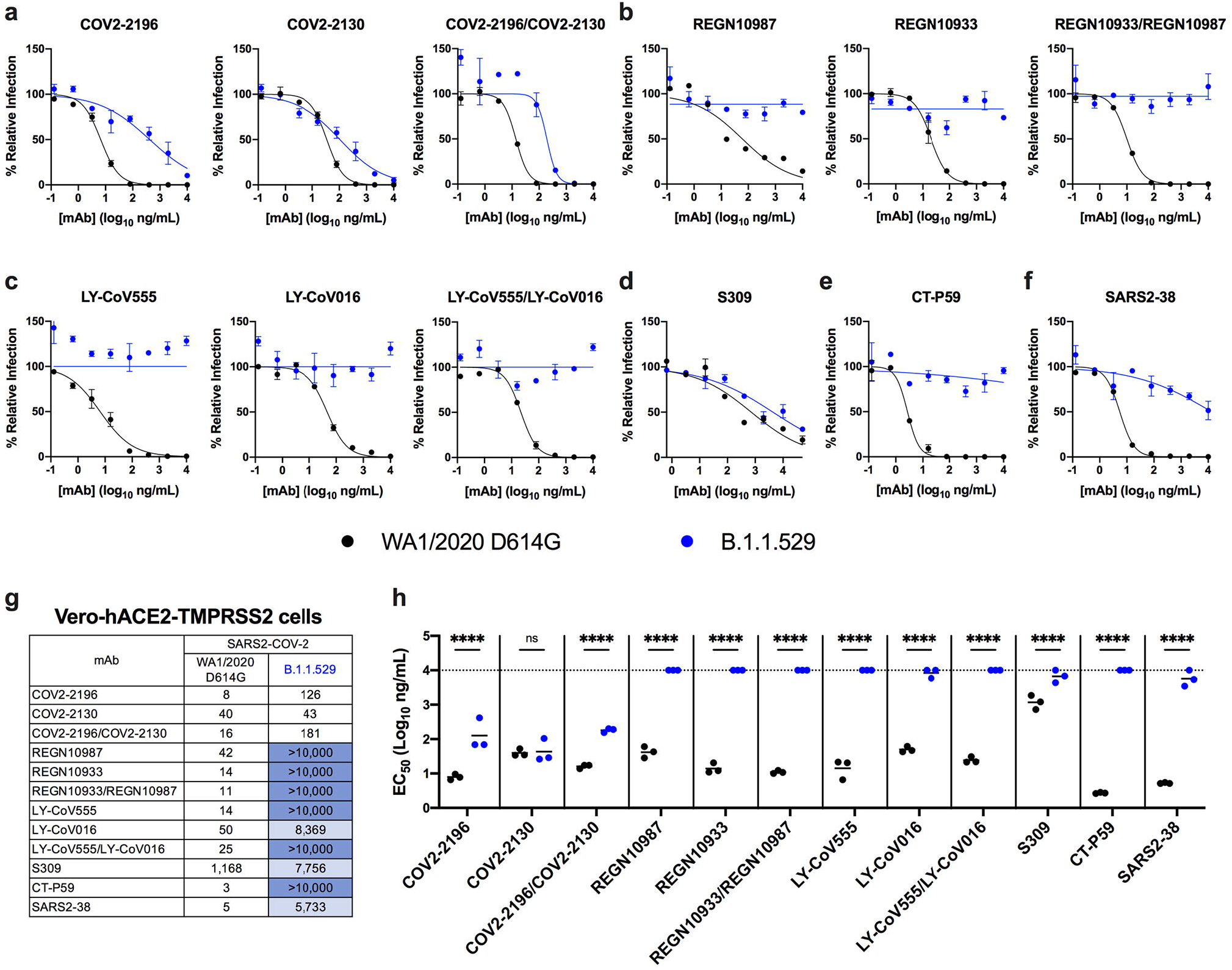
Neutralization of SARS-CoV-2 B.1.1.529 Omicron strain by mAbs in Vero-hACE2-TMPRSS2 cells. **a-f**, Neutralization curves in Vero-hACE2-TMPRSS2 cells comparing the sensitivity of SARS-CoV-2 strains with the indicated mAbs (S309, COV2-2196, COV2-2130; REGN10933, REGN10987, LY-CoV555, LY-CoV016, CT-P59, and SARS2-38) with WA1/2020 D614G and B.1.1.529. Also shown are the neutralization curves for antibody cocktails (COV2-2196/COV2-2130, REGN10933/REGN10987, or LY-CoV555/LY-CoV016). One representative experiment of three performed in technical duplicate is shown. Error bars indicate range. **g**, Summary of EC_50_ values (ng/ml) of neutralization of SARS-CoV-2 viruses (WA1/2020 D614G and B.1.1.529) performed in Vero-hACE2-TMPRSS2 cells. Data is the geometric mean of 3 experiments. Blue shading: light, EC_50_ > 5,000 ng/mL; dark, EC_50_ > 10,000 ng/mL. **h**, Comparison of EC_50_ values by mAbs against WA1/2020 D614G and B.1.1.529 (3 experiments, ns, not significant; ^****^, *P* < 0.0001; two-way ANOVA with Sidak’s post-test). Bars indicate mean values. The dotted line indicates the upper limit of dosing of the assay.

Our experiments show a marked loss of inhibitory activity by several of the most highly neutralizing mAbs that are in advanced clinical development or have EUA approval. We evaluated antibodies that correspond to monotherapy or combination therapy that have shown pre- and post-exposure success in clinical trials and patients infected with historical SARS-CoV-2 isolates. Our results confirm *in silico* predictions of how amino acid changes in B.1.1.529 RBD might negatively impact neutralizing antibody interactions^16,25^. Moreover, they agree with preliminary studies showing that several clinically used antibodies lose neutralizing activity against B.1.1.529 spike-expressing recombinant lentiviral or vesicular stomatitis virus (VSV)-based pseudoviruses^26-28^. One difference is that our study with authentic B.1.1.529 showed only moderately reduced neutralization by antibodies corresponding to the AstraZeneca combination (COV2-2196 and COV2-2130); in contrast, another group reported escape of these mAbs using a VSV pseudovirus displaying a B.1.1.529 spike protein in Huh7 hepatoma cells^27^. Additional studies are needed to determine whether this disparity in results is due to the cell type, the virus (authentic versus pseudotype), or preparation and combination of antibody.

While the Regeneron (REGN10933 and REGN10987), Lilly (LY-CoV555 and LV-CoV016) and Celltrion (CT-P59) antibodies or combinations showed an almost complete loss of neutralizing activity against B.1.1.529, in our assays with Vero-TMPRSS2 and Vero-hACE2-TMPRSS2 cells, the mAbs corresponding to the AstraZeneca combination (COV-2196 and COV-2130) or Vir Biotechnology (S309) products retained substantial inhibitory activity. Although these data suggest that some of mAbs in clinical use may retain benefit, validation experiments *in vivo*^7^ are needed to support this conclusion and inform clinical decisions.

Given the loss of inhibitory activity against B.1.1.529 of many highly neutralizing anti-RBD mAbs in our study, it appears likely that serum polyclonal responses generated after vaccination or natural infection also may lose substantial inhibitory activity against B.1.1.529, which could compromise protective immunity and explain a rise in symptomatic infections in vaccinated individuals^29^. Indeed, studies have reported approximately 25 to 40-fold reductions in serum neutralizing activity compared to historical D614G-containing strains from individuals immunized with the Pfizer BNT162b2 and AstraZeneca AZD1222 vaccines^26,28,30,31^.

We note several limitations of our study: (1) Our experiments focused on the impact of the extensive sequence changes in the B.1.1.529 spike protein on mAb neutralization in cell culture. Despite observing differences in neutralizing activity with certain mAbs, it remains to be determined how this finding translates into effects on clinical protection against B.1.1.529; (2) Although virus neutralization is a correlate of immune protection against SARS-CoV-2^7,32,33^, this measurement does not account for Fc effector functions if antibodies residually bind B.1.1.529 spike proteins on the virion or surface of infected cells. Fcγ receptor or complement protein engagement by spike binding antibodies could confer substantial protection^34-36^; (3) We used the prevailing B.1.1.529 Omicron isolate that lacks an R346K mutation. While only 8.3% of B.1.1.529 sequences in GISAID (accessed on 12/14/2021) have an R346K mutation, this substitution might negatively impact neutralization of some EUA mAbs given that it is a crystallographic contact for COV2-2130, REGN10987, and S309 (**Fig 1l**). At least for S309, the R346K mutation did not impact neutralization of pseduoviruses displaying B.1.1.529 spike proteins^28^. Nonetheless, studies with infectious B.1.1.529 isolates with R346K mutations may be warranted if the substitution becomes more prevalent; (4) Our data is derived from experiments with Vero-TMPRRS2 and Vero-hACE2-TMPRSS2 cells. While these cells standardly are used to measure antibody neutralization of SARS-CoV-2 strains, primary cells targeted by SARS-CoV-2 *in vivo* can express unique sets of attachment and entry factors^37^, which could impact receptor and entry blockade by specific antibodies. Indeed, prior studies have reported that the cell line used can affect the potency of antibody neutralization against different SARS-CoV-2 variants^6^.

In summary, our cell culture-based analysis of neutralizing mAb activity against an authentic infectious B.1.1.529 Omicron SARS-CoV-2 isolate suggests that several, but not all, existing therapeutic antibodies will lose protective benefit. Thus, the continued identification and use of broadly and potently neutralizing mAbs that target the most highly conserved residues on the SARS-CoV-2 spike likely is needed to prevent resistance against B.1.1.529 and future variants with highly mutated spike sequences.

## ACKNOWLEDGEMENTS

This study was supported by grants and contracts from NIH (R01 AI157155, U01 AI151810, 75N93021C00014, HHSN272201700060C, and 75N93019C00051), the Defense Advanced Research Project Agency (HR0011-18-2-0001), the Japan Program for Infectious Diseases Research and Infrastructure (JP21wm0125002) from the Japan Agency for Medical Research and Development (AMED), and the Dolly Parton COVID-19 Research Fund at Vanderbilt University Medical Center. We thank Rachel Nargi, Robert Carnahan, Tiong Tan, and Lisa Schimanski for assistance and generosity with mAb generation and purification, and Samuel A. Turner from the Center for Pathogen Evolution at the University of Cambridge for evaluating B.1.1.529 sequences.

## AUTHOR CONTRIBUTIONS

L.A.V. performed and analyzed neutralization assays. P.J.H. and L.A.V. propagated SARS-CoV-2 viruses. P.H. performed sequencing analysis. J.E.C., S.J.Z., L.P., and D.C. generated and provided mAbs. J.M.E. and D.H.F. performed structural analysis. J.E.C., Y.K., and M.S.D. obtained funding and supervised the research. L.A.V. and M.S.D. wrote the initial draft, with all other authors providing editorial comments.

## COMPETING FINANCIAL INTERESTS

M.S.D. is a consultant for Inbios, Vir Biotechnology, Senda Biosciences, and Carnival Corporation, and on the Scientific Advisory Boards of Moderna and Immunome. The Diamond laboratory has received funding support in sponsored research agreements from Moderna, Vir Biotechnology, and Emergent BioSolutions. J.E.C. has served as a consultant for Luna Biologics and Merck Sharp & Dohme Corp., is a member of the Scientific Advisory Boards of Meissa Vaccines and is Founder of IDBiologics. The Crowe laboratory has received sponsored research agreements from Takeda Vaccines, AstraZeneca and IDBiologics. Vanderbilt University has applied for patents related to two antibodies in this paper. L.A.P. and D.C. are employees of Vir Biotechnology and may hold equity in Vir Biotechnology. L.A.P is a former employee and shareholder in Regeneron Pharmaceuticals.

## METHODS

### Cells

Vero-TMPRSS2^39^ and Vero-hACE2-TMPRRS2^6^ cells were cultured at 37°C in Dulbecco’s Modified Eagle medium (DMEM) supplemented with 10% fetal bovine serum (FBS), 10 mM HEPES pH 7.3, and 100 U/ml of penicillin–streptomycin. Vero-TMPRSS2 cells were supplemented with 5 µg/mL of blasticidin. Vero-hACE2-TMPRSS2 cells were supplemented with 10 µg/mL of puromycin. All cells routinely tested negative for mycoplasma using a PCR-based assay.

### Viruses

The WA1/2020 recombinant strain with substitutions (D614G) was described previously^40^. The B.1.1.529 isolate (hCoV-19/USA/WI-WSLH-221686/2021) was obtained from a midturbinate nasal swab and passaged once on Vero-TMPRSS2 cells as described^41^. All viruses were subjected to next-generation sequencing (GISAID: EPI_ISL_7263803) to confirm the stability of substitutions. All virus experiments were performed in an approved biosafety level 3 (BSL-3) facility.

### Monoclonal antibody purification

The mAbs used in this paper (COV2-2196, COV2-2130, S309, REGN10933, REGN10987, LY-CoV555, LY-CoV016, CT-P59, SARS2-38) have been described previously^12,15,19,42-46^. COV2-2196 and COV2-2130 mAbs were produced after transient transfection using the Gibco ExpiCHO Expression System (ThermoFisher Scientific) following the manufacturer’s protocol. Culture supernatants were purified using HiTrap MabSelect SuRe columns (Cytiva, formerly GE Healthcare Life Sciences) on an AKTA Pure chromatographer (GE Healthcare Life Sciences). Purified mAbs were buffer-exchanged into PBS, concentrated using Amicon Ultra-4 50-kDa centrifugal filter units (Millipore Sigma) and stored at −80□°C until use. Purified mAbs were tested for endotoxin levels (found to be less than 30 EU per mg IgG). Endotoxin testing was performed using the PTS201F cartridge (Charles River), with a sensitivity range from 10 to 0.1 EU per mL, and an Endosafe Nexgen-MCS instrument (Charles River). S309, REGN10933, REGN10987, LY-CoV016, LY-CoV555, CT-P59, and SARS2-38 mAb proteins were produced in CHOEXPI or EXPI293F cells and affinity purified using HiTrap Protein A columns (GE Healthcare, HiTrap mAb select Xtra #28-4082-61). Purified mAbs were suspended into 20 mM histidine, 8% sucrose, pH 6.0 or PBS. The final products were sterilized by filtration through 0.22 μm filters and stored at 4°C.

### Focus reduction neutralization test

Serial dilutions of mAbs were incubated with 10^2^ focus-forming units (FFU) of SARS-CoV-2 (WA1/2020 D614G or B.1.1.529) for 1 h at 37°C. Antibody-virus complexes were added to Vero-TMPRSS2 or Vero-hACE2-TMPRSS2 cell monolayers in 96-well plates and incubated at 37°C for 1 h. Subsequently, cells were overlaid with 1% (w/v) methylcellulose in MEM. Plates were harvested at 30 h (WA1/2020 D614G on Vero-TMPRSS2 cells), 70 h (B.1.1.529 on Vero-TMPRSS2 cells), or 24 h (both viruses on Vero-hACE2-TMPRSS2 cells) later by removal of overlays and fixation with 4% PFA in PBS for 20 min at room temperature. Plates with WA1/2020 D614G were washed and sequentially incubated with an oligoclonal pool of SARS2-2, SARS2-11, SARS2-16, SARS2-31, SARS2-38, SARS2-57, and SARS2-71^47^ anti-S antibodies. Plates with B.1.1.529 were additionally incubated with a pool of mAbs that cross-react with SARS-CoV-1 and bind a CR3022-competing epitope on the RBD^19^. All plates were subsequently stained with HRP-conjugated goat anti-mouse IgG (Sigma, A8924) in PBS supplemented with 0.1% saponin and 0.1% bovine serum albumin. SARS-CoV-2-infected cell foci were visualized using TrueBlue peroxidase substrate (KPL) and quantitated on an ImmunoSpot microanalyzer (Cellular Technologies). Antibody-dose response curves were analyzed using non-linear regression analysis with a variable slope (GraphPad Software), and the half-maximal inhibitory concentration (EC_50_) was calculated.

### Model of mAb-B.1.1.529 spike complexes

The spike model is a composite of data from PDB: 7C2L and PDB: 6W41. Models of mAb complexes were generated from their respective PDB files with the following accession codes: COV2-2196 (PDB: 7L7D); COV2-2130 (PDB: 7L7E); S309 (PDB: 6WPS); REGN-10987 (PDB: 6XDG); REGN-10933 (PDB: 6XDG)); LY-CoV555 (PDB: 7KMG) LY-CoV016 (PDB: 7C01); CT-P59 (PDB: 7CM4) and SARS2-38 (PDB: 7MKM). Epitope footprints used in the multiple sequence alignment were determined using PISA interfacial analysis on the various mAb:RBD complexes^48^. Structural figures were generated using UCSF ChimeraX^49^.

## Data availability

All data supporting the findings of this study are available within the paper and are available from the corresponding author upon request.

## Code availability

No code was used in the course of the data acquisition or analysis.

## Reagent availability

All reagents described in this paper are available through Material Transfer Agreements.

## Statistical analysis

The number of independent experiments and technical replicates used are indicated in the relevant Figure legends. A two-way ANOVA with Sidak’s post-test was used for comparisons of antibody potency between WA1/2020 D614G and B.1.1.59.

## Notes

### Competing Interest Statement

M.S.D. is a consultant for Inbios, Vir Biotechnology, Senda Bioscences, and Carnival Corporation, and on the Scientific Advisory Boards of Moderna and Immune. The Diamond laboratory has received funding from Moderna, Vir Biotechnology, and Emergent BioSolutions. J.E.C. has served as a consultant for Luna Biologics and Merck Sharp and Dohme Corp., is a member of the Scientific Advisory Boards of Meissa Vaccines, and is Founder of IDBiologics. The Crowe laboratory has received sponsored research agreements from Takeda Vaccines, AstraZeneca and IDBiologics. Vanderbilt University has applied for patents related to two antibodies in this paper. L.A.P. and D.C. are employees of Vir Biotechnology and may hold equity in Vir Biotechnology. L.A.P is a former employee and shareholder in Regeneron Pharmaceuticals.

